# Utilization and prospectus of non-timber forest products as livelihood materials

**DOI:** 10.1101/2020.10.19.345223

**Authors:** Atanu Kumar Das, Md. Asaduzzaman, Md Nazrul Islam

## Abstract

A study was conducted to find out the present utilization of non-timber forest products of the Sundarbans and tries to find out the alternative uses of these resources. The Sundarbans is the largest mangrove forest in the world and proper utilization of all resources can get a chance to manage this forest in sustainable way. A questionnaire survey was conducted to get information about the utilization of non-timber forest products. It has been found that about 87% of the people are fully dependent and 13% of the people are partially dependent on the Sundarbans. Among minor forest products, golpata, honey and fish were used by 92%, 93% and 82% of people, respectively. Most of the people are unknown about the alternative use of minor forest products but there is a great chance to use them for better purposes. Alternative uses of these products will help to improve the forest conditions as well as the socio-economic conditions of the people adjacent to the Sundarbans.

## Introduction

Non-Timber Forest Products (NTFPs) are biological origin derived from the forest, other wooded land and trees outside the forest (FAO, 1999). According to Shiva and Mathur (2007) Minor Forest Products (MFP) or Non-Wood Forest Products (NWFPs) or Non-Timber Forest Products (NTFPs) are all products obtained from plants of forest origin and host plant species yielding products in association with insect and animals or they are parts and items of mineral origin except timber. The importance of these forest products makes it imperative to employ a sustainable management mechanism for the rapidly depleted forest resources to maintain an uninterrupted supply of these resources for the future generation. Sustainable development implies development, which while protecting the environment, allows a type of economic activity that can be sustainable into the future with minimum damage to people or ecosystem (Goudie, 2000). These products are called as minor forest products, because of inadequate documentation of their trade in international markets. However, with appropriate market development, foreign exchange can be earned from these minor forest products (Osemeobo and Ujor, 1999).

The Sundarbans, world’s largest mangrove forest, is located at the southern extremity of the Ganges river delta bordering on the Bay of Bengal (Hussain and Karim, 1994). The forest covers and area of about 10,000 m^2^ of which 62% falls within the territory of Bangladesh while the remaining area belongs to India (Siddiqi, 2001). The Bangladesh Sundarbans lies between the latitudes 21°31’ - 22°30’ N and between the longitudes 89°00’ - 90°00’E. About 4,143 sq. km of a total 6,017 sq. km is landmass and the remaining 1,874 sq. km are water bodies in the form of numerous rivers, canals and creeks of widths varying from a few meters to several kilometers (Chaffey *et al.*, 1985). The Sundarbans supports rich and diverse non-wood produce, many of which are indispensable to the adjoining rural people (Siddique, 2001). Uttar Bedkashi and Koyra Sadar (1 No. Koyra) are the two union parishad of Koyra upazila of Khulna district 7 which are adjoining to the Sundarbans. A substantial amount of revenue of the government is earned through non-timber forest products. The resource is of both plant and animal origin. Plant based non-wood produce includes golpata (*Nypa fruticans*), hantal (*Phoenix paladosa*), grasses (Nal, Ulu, Hogla, Malia), medicinal plants etc. As regards animals, income is generated from scale of fish, crabs, shells, shrimp fry, honey, wax etc. Non-timber provides fuelwood, thatching material, house post, medicine, food and material for cottage industries and domestic use of the surrounding people.

The limited resources of the forest are being indiscriminately extracted due to dependency on forest resources for their lively hood. Besides, people are not adequately informed about the resources of the forest, how to use the resources of the forest and other pertaining information. Moreover, the people directly involved with the Sundarbans for their livelihood are not being effectively addressed by the government organizations. In this study, it was conducted to identify the present uses of non-timber forest products and to identify the alternative uses of the non-timber forest products of the Sundarbans for better uses for the socio-economic development of the people adjacent to the Sundarbans ultimately for the country.

## Materials and Methods

The study was conducted in Uttar Bedkashi and Koyra Sadar Union of Koyra Upziala of Khulna District, purposively selected to investigate the situation of household utilization of minor forest products of the Sundarbans. The study area comprises two union named Uttar Bedkasi and Koyra Sadar of Koyra upazilla (Khulna district). Paikgachha upazilla on the north, the Bay of Bengal and the Sunderban bound Koyra upazilla on the south, Dacope upazilla on the east, Assasuni and Shyamnagar upazilla on the west (Banglapedia, 2008).

A semi-structural questionnaire was prepared keeping the objectives of the study in view. The schedule was pre-tested before final collection of data. The study was conducted followed by the Quota (Non-probability) Sampling method. Data was collected by the questionnaire survey of the sample respondents. Direct interview method was followed. The secondary data was collected from web sites, Seminar library of Forestry and Wood Technology Discipline, Khulna University.

After collecting information from primary and secondary sources, data were processed by the three steps i.e. Step-1: Reviewed of collected data and information, Step-2: Discarded of unnecessary part of the information and data and Step-3: Stored of revised data and information. MS Excel 2007 package was used for the purpose of data analysis.

## Results and Discussions

### Socio-economic condition

Socio-economic status has been presented in Table 1. Majority of the respondents(78%) belonged to young to middle age category having 60% primary to secondary level of education and 36% was illiterate with 93% had small to medium family size. 74 percent of the respondents’ family member go for education and their education level is primary to secondary. Majority of the respondents (73%) belonged to marginal to medium land holder and 6% were land less. 65 percent family posse’s hen and 40 percents family has boat. 95 percent respondents were engaged in fishing with 99 percent were the male as earning member. Low income group was 71 percent and 92 percent respondents’ expenditure was higher than income. Over expenditure family always owe to different types of NGO for lending money.

**Table 1:** Salient features of the respondents

| Characters | Observed range of score | Categories | Percentage of Respondents |
| --- | --- | --- | --- |
| Age (years) | 19 to 70 | Young aged (Up to 30) | 30 |
|  |  | Middle aged (31 to 45) | 48 |
|  |  | Old aged (Above 46) | 22 |
| Level of education (Schooling years) | 0 to above higher secondary | Illiterate(No schooling) | 36 |
|  |  | Primary (1-5) | 40 |
|  |  | Secondary (6-10) | 20 |
|  |  | Higher secondary (11-12) | 1 |
|  |  | Above higher secondary (>12) | 3 |
| Family size (No.) | 3 - 11 | Small (up to 6) | 64 |
|  |  | Medium (7-9) | 29 |
|  |  | Large (>10) | 7 |
| Level of Family Education(schooling years) no. of family | Primary to Higher degree | Primary | 47 |
|  |  | Secondary | 27 |
|  |  | Higher secondary | 5 |
|  |  | Higher degree | 2 |
| Total Land(decimal) | Land less to 363 decimal | Land less | 6 |
|  |  | Marginal(up to 10 decimal) | 41 |
|  |  | Medium (11decimal to 60 decimal ) | 32 |
|  |  | Large (above 60 decimal) | 21 |
| Other assets (Possessing no. of family) |  | Livestock |  |
|  |  | Cow | 11 |
|  |  | Sheep | 3 |
|  |  | Goat | 44 |
|  |  | Hen | 65 |
|  |  | Duck | 41 |
|  |  | Radio | 14 |
|  |  | Tape recorder | 4 |
|  |  | TV | 14 |
|  |  | Boat | 40 |
|  |  | Net | 5 |
|  |  | Mobile | 21 |
| Types of earning member | Child to female | Child as supporting | 9 |
|  |  | Female as head of the family | 1 |
|  |  | Female as supporting | 12 |
|  |  | Male | 99 |
|  |  | Fisher man | 95 |
|  |  |  | 2 |
| Profession |  | Doctor | 1 |
|  |  | Farmer | 1 |
|  |  | Carpenter | 1 |
|  |  | Van driver |  |
| Monthly Income (Tk.) | 1200 to 10,000 | Low (Up to Tk. 3000) | 71 |
|  |  | Medium (3100-6000) | 24 |
|  |  | High (Above 6000) | 5 |
| Monthly Expenditure (Tk.) |  | Over income | 92 |
|  |  | Below income | 5 |
|  |  | Equal to income | 3 |

### Utilization and prospectus of non-timber forest products

Dependency on non-timber forest products cannot be denied by any one because they use fish at least. In this study, we have got same type of information during our survey; however, 87% are fully dependent on the Sunderbans for non-timber forest products. Majority of the people (87%) are involved directly to collect minor forest products as they live near by the Sundarbans. Their main income source is also the Sunderbans. People are poor and they live hand to mouth. Most of the people have no own boat, net and they also face economical problem. Though they have thousands of problems, they (97%) collect their non-timber forest product as ownership by sharing process.

There are many minor forest products of the Sundarbans such as golpata(*Nypa fruticans*), honey and bee’s wax, hantal(*Phoenix paladosa*), fish, shrimp fry, medicinal plants, fuelwood. Uses, percentage, availability and price of minor forest products have been presented in Table 2.

**Table 2.**
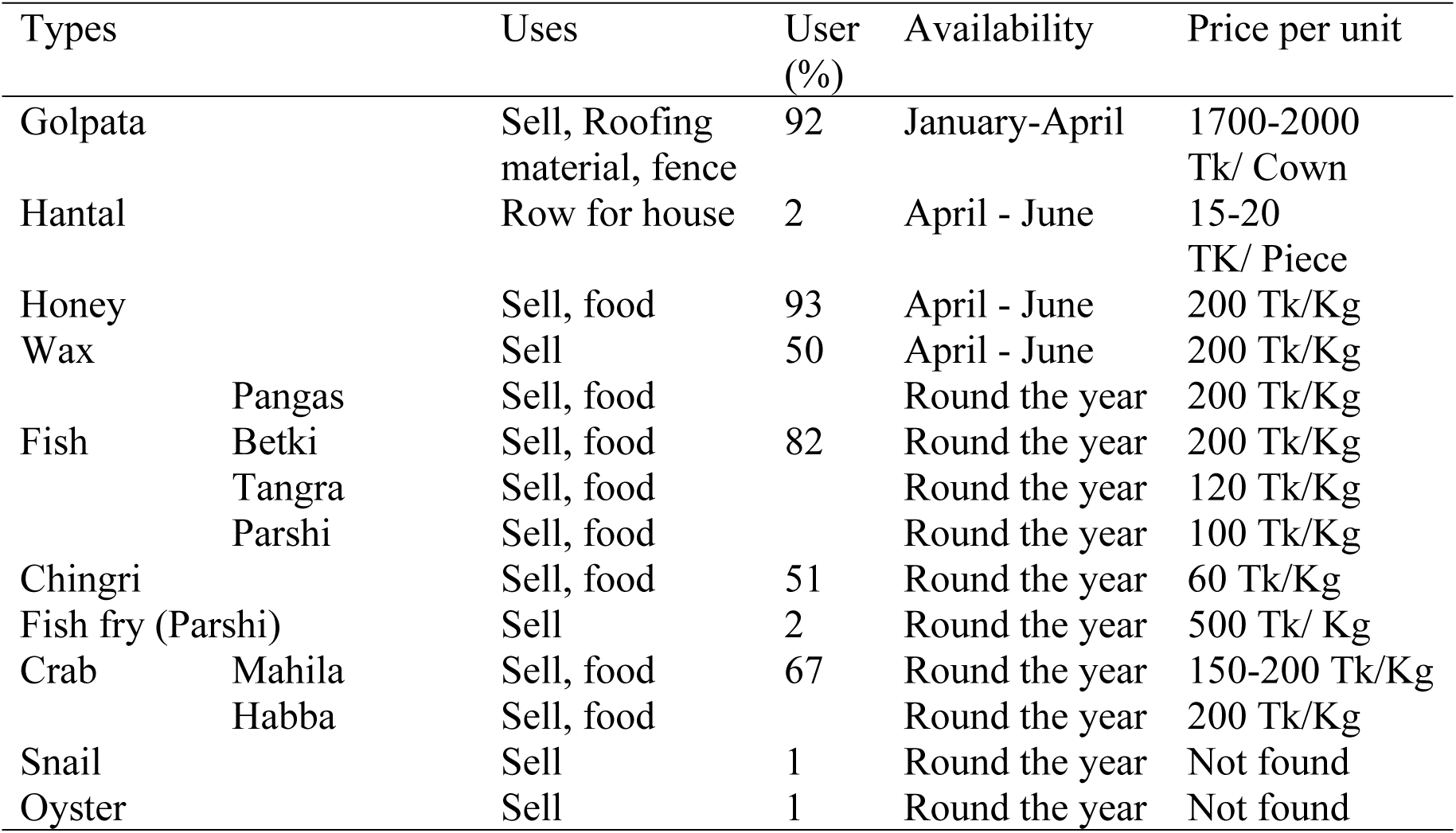
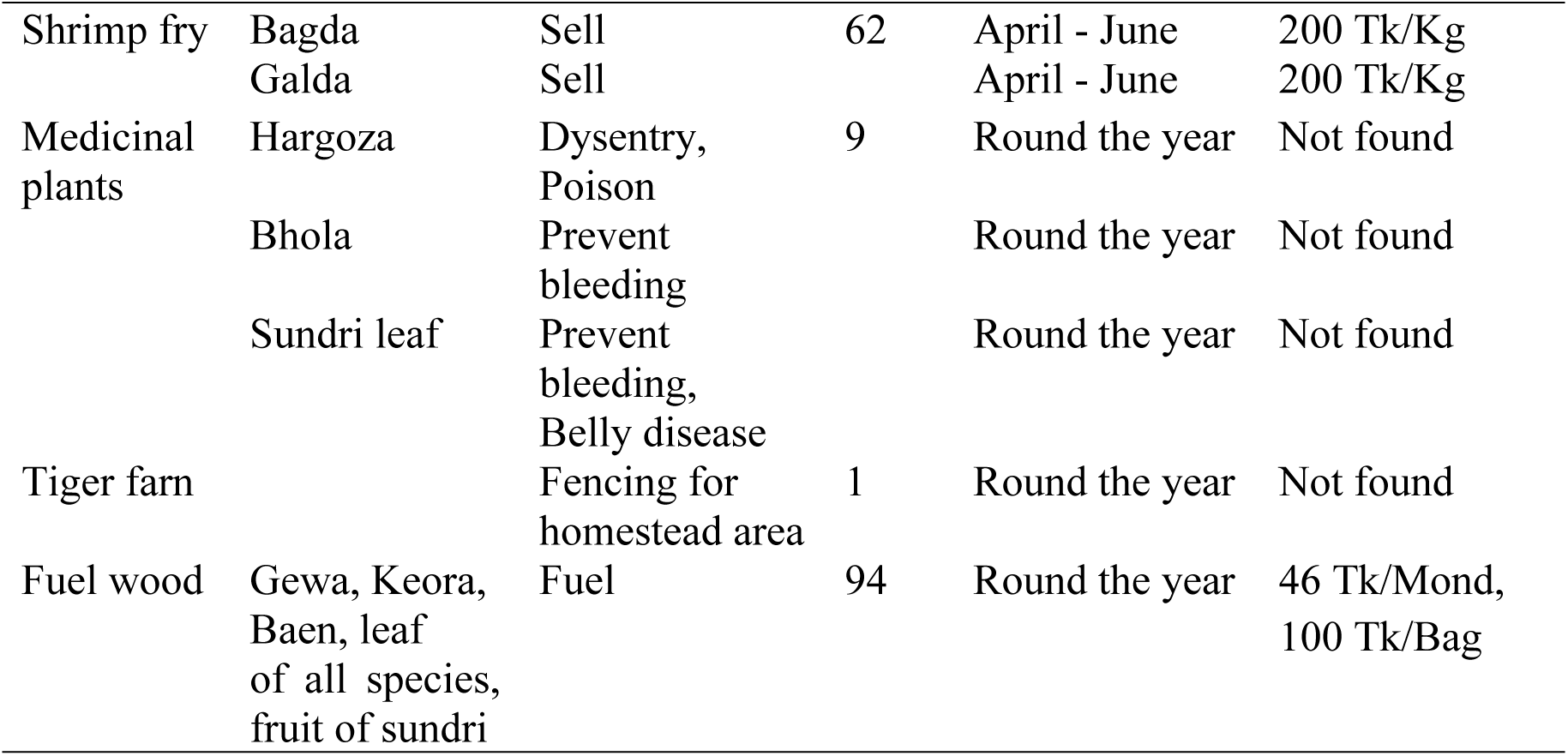
Types and uses of minor forest products

In our study area, we have found that golpata (*Nypa fruticans*) is used it as roofing material and 92% people use it for this purpose. They also sell it in the market to earn money. Local people prepare sweet from the fruit of golpata. On the other hand, hantal (*Phoenix paladosa*) is used as row of house and only 1% people use it. Tiger farn (*Acrostichum aureum*) or hudo is used as fence of homestead but only 1% has been seen to use it for this purpose. Some mangrove species have medicinal value. Bhola (*Hibiscus tiliaceus*), horgoza (*Acanthus olicifolius*), sundri (*Heritiera fomes*) are one of them to use as medicine in our study area. Bhola is used for dysentery. There are only 9% people use medicinal plants of the Sundarbans in our study area. Leaf of all species and fruit of sundri are the main source of fuelwood of this area. People collect it from the river. Beside this, gewa, keora, baen are also used as fuelwood directly. The user of fuel wood percentage is 94.

Honey collectors mainly sell honey and they also use it as food. Others people buy it for food and medicine. Honey and wax is used 93% and 50% people respectively of this study area. Shrimp fry is particularly important income source of the fisherman in this study area. They collect it for selling to live on and 62% people use it. The local names of these fry are bagda and golda. The people of the study area also catch parshi fry and they sell it to different fish farm. Only 2% people use it to earn by selling in the market. On the other hand, the Sundarbans is full of fish resources and many fish species are available in this area. Among many species betki, pangas, tangra, parsi etc. are remarkable. People sell this as well as use it as food. The dependency on fish is 82% in this area. People of this area earn by selling crab and this type of people is 33% but 51% people use chingri to sell for earning money. Snail and Oyster are the two important sources shell and these are mainly used to producing lime. Very few people are engaged for this purpose in this area. Only 1% people use it to earn money.

The availability of non-timber forest products vary with products. Tiger farn, medicinal plants, chngri, fish fry (parsi), crab, snail and oyster are available round the year in the study area. Golpata is available from January to April whether hantal, honey, wax and shrimp fry are available from April to June. Most of the non-timber forest products are sold as Tk/kg and the range of price is 150 −200 Tk/kg. Golpata is sold as 1700 - 2000 Tk/Cown but fuelwood is sold as 46 Tk/Mond or 100 Tk/Bag.

There is a great scope for launching cottage industry in the study area because of presence of some species. Hantal can be possible to produce house hold furniture. On the other hand, candle can be possible to produce from wax; mat and cap can be possible to produce from some grasses. Sweet can also be possible to produce from the juice of golpata. These type of activities will help to improve the socio-economic condition of the study area ultimately of Bangladesh.

## Conclusion

Major portion of the people of Koyra Sadar and Uttar Bedkashi Union are dependent on minor forest products of the Sundarbans. Even government also earns a large amount of revenue from the non-timber forest products. Alternative uses of these non-timber forest products will help to earn more for government and the local people adjacent Sunderbans. This will also help to improve the life style of the people adjacent to the Sundebans. Therefore, the government should come forward first to take care about this important forest products so that maximum benefit can be obtained from the Sundarbans. Other non-government organization should also take part to help the government to manage it efficiently.

## Acknowledgement

The authors would like to thank the local people of the study area to complete this type of research smoothly.

